# Adaptive Clonal Expansion Shapes Brain Development

**DOI:** 10.1101/2025.03.04.641452

**Authors:** Giulia Di Muzio, Sarah Benedetto, Li-Chin Wang, Lea Weber, Franciscus van der Hoeven, Brittney Armstrong, Hsin-Jui Lu, Jana Berlanda, Verena Körber, Nina Claudino, Michelle Krogemann, Thomas Höfer, Pei-Chi Wei

## Abstract

Embryonic neural stem/progenitor cells (NSPCs) exhibit remarkable proliferative plasticity, allowing them to fully recover neuronal populations even after substantial cell loss ^1,2^. However, it remains unclear whether all embryonic NSPCs respond to brain lesions. To address this, we developed a mouse model to investigate NSPC proliferation dynamics, hypothesizing that the loss of progenitor cells would induce fitness competition among NSPCs. In this model, half of the founder NSPCs were ablated using diphtheria toxin A at the onset of neurogenesis, yet the surviving cells regenerated a brain containing all neuronal types within five of the total twenty embryonic days. Analysis of allelic variants revealed overrepresented somatic variants, indicating that only a small fraction of NSPCs underwent significant clonal expansion during early neurogenesis. Modelling proliferation dynamics predicted that as few as 10% of NSPCs could produce 83% of neurons by the time of birth. Single nucleotide substitution analysis suggested a potential link to oxidative metabolism in some of the expanded clones. Moreover, single-cell transcriptomics showed delayed development and a reduced NSPC pool as consequences of adaptive clonal expansion. Our findings highlight that NSPC exhibit varying expansion potential and that adaptive clonal expansion indirectly altered neuronal cell composition in the brain.

## Background

The proliferation of neural stem/progenitor cells (NSPCs) is crucial for laying the foundation of mammalian brain development. This process generates a large pool of neural progenitors that will differentiate into neurons and glial cells in the brain. It also shapes the basic structure and size of the developing brain^3,4^. Moreover, the early expansion of NSPCs influences the final size of specific brain regions^5,6^. In certain species, this expansion contributes to the formation of cortical folds, which increase the cortical surface area within the confined space of the skull ^3,7^.

The proliferation dynamics of NSPCs directly determine the number of progenies each cell produces. As NSPCs replicate their DNA during each proliferation cycle, all progenies within the same neuronal clone inherit the DNA blueprint as their cell of origin. Errors in this blueprint can be passed down to neurons, potentially contributing to neurological disorders ^8,9^. For example, in some cases, errors present in only 1% of neurons are sufficient to cause focal cortical dysplasia^10^. Therefore, investigating the expansion capacity of NSPCs is critical for understanding how erroneous genetic blueprints are distributed across different brain regions.

Recent studies on the neuronal clone size in the human brain have revealed that approximately 90-200 neural progenitor cells are present at the time of midline establishment^11^. In hindsight, one might expect an equal representation of neuronal clones derived from each individual founder NSPC. However, somatic variants-dependent lineage-tracing studies suggested otherwise. In up to a quarter of human brains, neuronal clones comprising, bias in proportion, for more than 8% of neurons were found^12^. Additionally, a study analysing over 130 human brains has suggested evidence for selective expansion of neuronal progenitors in normal human brains^13^. These findings indicate that NSPC possess intrinsic capacities to generate a larger number of neurons. However, within the tightly constrained period of embryonic neurogenesis, the extent to which this potential can be exploited during early neurogenesis remains unknown.

Although it has been speculated that NSPCs exhibit differential proliferative abilities^14,15^, answering this question remains challenging. Existing experiment approaches have either focused on collective population-level proliferative behavior^14,15^ or imposed all-or-none conditions^16–19^, making it difficult to capture the heterogeneous nature of NSPC proliferation. Additionally, while alterations in the proliferation dynamics of neural progenitor cells can directly impact the spread of brain-specific mutations, it remains unclear whether clonal expansion dynamics can exceed pathological thresholds and what risk factors may trigger unbalanced clonal expansion. Therefore, an animal model that allows NSPCs to exhibit their natural proliferative dynamics is needed.

We present a mouse model in which a reduction in NSPCs at the onset of neurogenesis promotes selective clonal expansion. We analysed *in vivo* changes in cell proliferation rates and examined neuronal differentiation status at the end of embryonic neurogenesis. We used whole-genome sequencing to evaluate clonal expansion, by accessing variant allele frequency distributions to infer the clonal population size. Our analysis revealed the presence of dominant clones, with the largest comprising 50% of neuronal cells, contributing significantly to the overall neuronal population. Furthermore, we simulated clonal expansion dynamics throughout embryonic neurodevelopment using a pool-size-weighted mathematical model. Finally, through transcription factor activity profiling and co-regulated gene expression analysis in single cells, we concluded that adaptive clonal expansion reflects a developmental crisis, potentially posing risks to brain health.

## Results

### Creating a mouse model allowing NSPCs exert their expansion potential

The ability of the NSPCs to survive and proliferate can be influenced by stochastic developmental bottlenecks, such as high frequency of programmed cell death^20^, where unfit cells were eliminated through programmed cell death^17,21^. We hypothesize that embryonic cell death in brain creates a development bottleneck, encouraging NSPCs to expand.

To test this hypothesis, we created chimeric mice using host blastocysts encoding for the attenuated diphtheria toxin A (DTA)-expression cassette at the *ROSA26* locus (*R26-DTA*). The DTA expression was controlled by removing the 5’ end floxed stop cassette by the Cre recombinase, which was controlled by a transgenic rat *Nestin* promoter (*NesCre*)^22,23^. Prior research has shown that the rat *Nestin* promoter only activates in NSPCs - radial glial (RG) cells and the intermediate progenitor (IP) cells - at embryonic day (E) 12.5, but not at time points before the emergence of RG cells^22^. In addition, although DTA leakiness was known^24^, it did not affect our ability to address the expansion capacity of surviving NSPCs. Thus, selectively eliminating NSPCs by internally expressing DTA, represents an appropriate approach to address our question. In the following context, we refer to blastocysts containing *NesCre* and *R26-DTA* expression cassettes as *NesCre::R26-DTA* embryos (**Fig. 1A**). The control blastocysts lacking Cre recombinase expression are referred to as *R26-DTA* embryos.

**Figure 1.**
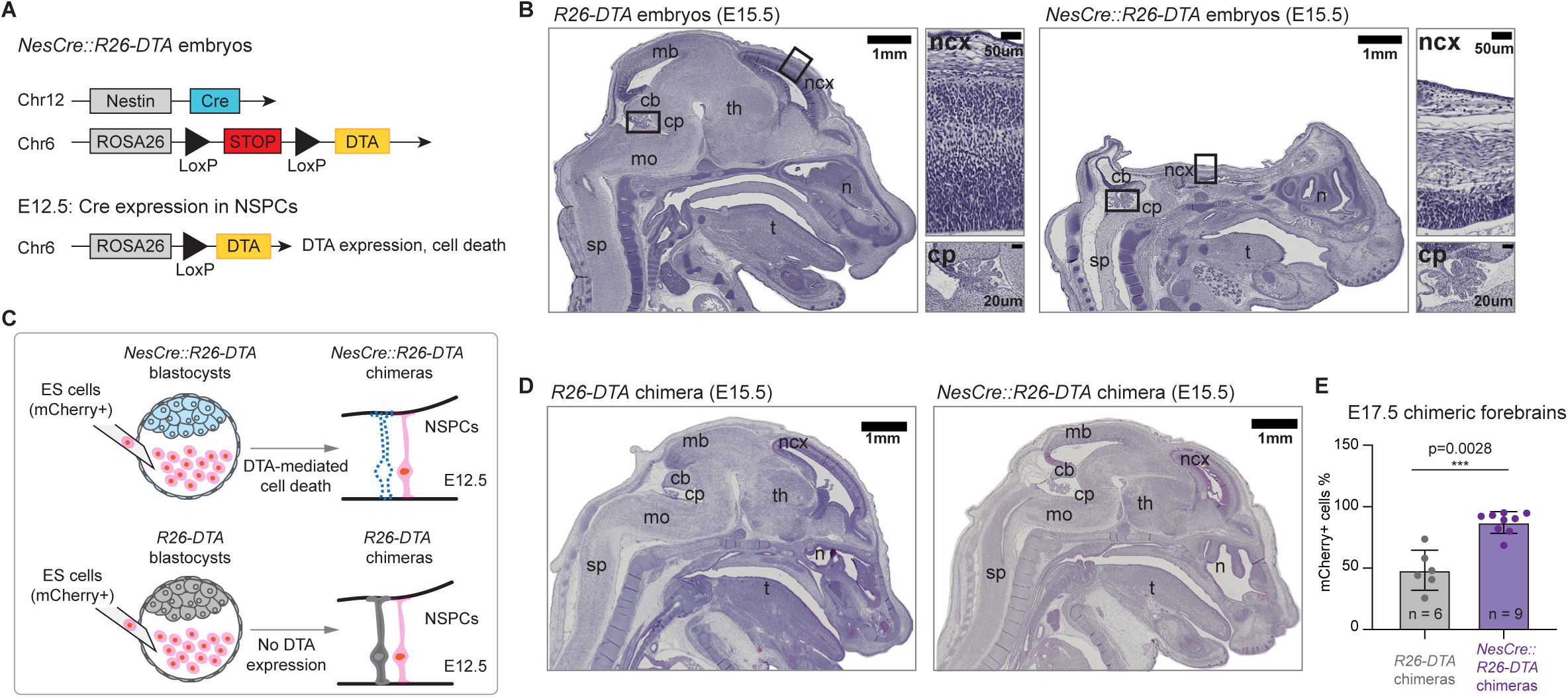
Creating the adaptive NSPC expansion model, the *NesCre::R26-DTA* chimeras. **(A)** Schematic representation of the transgenic expression cassettes in the genome of *NesCre::R26-DTA* embryos. Chr: chromosome. Nestin: rat transgenic Nestin promoter. Cre: Cre recombinase. ROSA26: ROSA26 locus on Chr6. DTA: diphtheria toxin subunit A. NSPCs: neural stem and progenitor cells. **(B)** A representation of hematoxylin and eosin (H&E) - stained sagittal brain sections of an *R26-DTA* embryo (left) and a *NesCre::R26-DTA* embryo (right) at embryonic day (E) 15.5. Insets showed higher magnification views of the rectangular areas of neocortex (ncx) or the choroid plexus (cp). Scale bars: 1 mm (overview), 50 µm (ncx view), and 20 µm (cp view). Brain structures, including midbrain (mb), neocortex (ncx), thalamus (th), medulla oblongata (mo), cerebellum (cb), and the spinal cord (sp) as well as craniofacial structures such as tongue (t), and nose cavity (n) were annotated. **(C)** Schematic of blastocyst complementation strategy. mCherry-positive embryonic stem cells (pink) were injected into host blastocysts (*R26-DTA* or *NesCre::R26-DTA*) to generate chimeras. The NSPCs in the *NesCre::R26-DTA* chimeras (dashed lined cell) died upon DTA expression, whereas the *R26-DTA* NSPCs (grey cell) were alive. **(D)** H&E-stained sagittal brain sections of an *R26-DTA* chimera (left) and a *NesCre::R26-DTA* chimera (right) at E15.5. **(E)** Barchart showed the proportion of mCherry+ cells in forebrain tissue of chimeras at E17.5. Data were presented as mean ± standard deviation, and the statistical significance was determined by unpaired two-tail Student t test. *** p < 0.005.

To benchmark DTA-induced NSPC death *in vivo*, we crossed male mice hemizygous for *NesCre* with female mice homozygous for the *R26-DTA* transgene (**Fig. S1A**). This approach produced about 50% *NesCre::R26-DTA* (n=16) and 50% control *R26-DTA* embryos (n=14), indicating that DTA expression did not affect embryonic mortality. Histological analysis revealed that the cerebral area was largely ablated in the *NesCre::R26-DTA* embryos at E12.5 (**Fig. S1B**) and E15.5 (**Fig. 1B**), confirming the failure of neurogenesis. Due to lack of brain tissue, the head structure of the *NesCre::R26-DTA* embryos collapsed during sample preparation for histology. In addition, immunofluorescent staining confirmed that a substantial lack of RG cells (PAX6-positive, **Fig. S1C,D**) and IP cells (TBR2-positive, **Fig. S1C,D**) in the brains of the E15.5 *NesCre::R26-DTA* embryos.

Next, to deplete part of the NSPCs, we created chimeric embryos containing a mixture of wild-type and *NesCre::R26-DTA* cells. This was achieved by blastocyst complementation, initially developed for immunology research^25^ and was applied for telencephalon development studies^1,2^. In brief, we injected wild-type mouse embryonic stem (ES) cells, which carry the constitutively expressed mCherry transgene in their genomes, into blastocysts to create *NesCre::R26-DTA* chimeras (**Fig. 1C**). In these chimeric embryos, half of the NSPCs were wild-type, while the other half carried the *NesCre::R26-DTA* transgene cassettes and subsequently died upon DTA expression. These embryos were referred to as *NesCre::R26- DTA* chimeras. In parallel, we injected wild-type mouse ES cells into *R26-DTA* blastocysts, creating *R26-DTA* chimeras as controls **(Fig. 1C)**.

We found that the injected mouse ES cells were incorporated into the central nervous system and reconstituted the brain and spinal cord tissue in the *NesCre::R26-DTA* chimeras (**Fig. 1D**). These data suggest that NSPCs in the *NesCre::R26-DTA* embryonic brains accelerate their proliferation rate, at least transiently. Given that our focus was on brain development, we did not investigate the spinal cord phenotype in this article. As expected, over 91 ± 5.3 % (mean ± SD) of forebrain cells were mCherry-positive in *Nestin-Cre::R26-DTA* chimeras, significantly higher than 48 ± 16.3 % (mean ± SD) mCherry-positive forebrain cells in *R26-DTA* chimeras (**Fig. 1E**). This bias was not due to differential ES cell contribution to the embryo itself, as the mCherry-positive cell proportion was comparable in tail fibroblasts isolated from *NesCre::R26-DTA* (67.5 ± 12.3%; mean ± SD) and *R26-DTA* chimeras (55.2 ± 11% %; mean ± SD) (**Fig. S1E**). We reasoned that the difference in the mCherry-positive ratio between tail fibroblasts and neuronal cells in the *Nestin-Cre::R26-DTA* chimeras resulted from complementation by the surviving wild-type NSPCs in the embryonic brains.

Active complementation also reconstituted all neuronal cell types at the end of embryonic neurogenesis. We analysed the gene expression for 61,080 (*NesCre::R26-DTA*) and 41,788 (*R26-DTA*) mCherry-expressing single cells isolated from the forebrain and the mid- and hindbrain (**Fig. S1F,G; Table S1**). In the *NesCre::R26-DTA* chimeras, the mCherry- positive cells represented all the neuronal cell types. We concluded that, in response to induced cell death, the living RG cells exerted their expansion potential to meet the demands of brain development.

### Selected clonal expansion in the *NesCre::R26-DTA* chimera’s brain

When NSPCs expands faster, they produce more progeny. This expansion can be driven either by all neuronal clones equally or by a specific subset of clones during early neurogenesis. Somatic mutations provide a way to trace neuronal clones, as any mutation that arises in the NSPC will propagate to all the neurons derived from it, allowing to associate specific DNA alterations to specific neuronal clones. To investigate which of the scenarios occurred during brain complementation, we performed deep whole-genome sequencing (WGS; mean coverage of 198x; **Table S2**) on the genomes of mCherry-positive cells isolated from the forebrain of three *R26-DTA* and six *NesCre::R26-DTA* chimeras at E17.5. We then analysed the frequency of small variants, specifically single nucleotide variants (SNVs) and small insertions and deletions (indels), acquired after the germ-layer separation at around E6.5^26^. To exclude variants present in the ES cell genomes or those that arose before germ-layer separation, we sequenced the genomes of mCherry-positive ES cells (55X coverage) and embryo-matched mCherry-positive tail fibroblasts (mean coverage of 76X; Table S2), which are derived from the mesoderm. Variants shared between tail fibroblasts and forebrains or ES cells and forebrains were removed for downstream analyses (**Fig. 2A**). Considering that only fifteen ES cells were injected into each blastocyst and about 70% of the ES cells contributed to the embryo proper^27,28^, we expect to identify variants that are specific to each ES cell at a minimum frequency of 10%. With deep sequencing coverage of forebrain cells, we expect to identify and remove ES cell-specific variants from the forebrain genomes, including those originating from culture artifacts.

**Figure 2.**
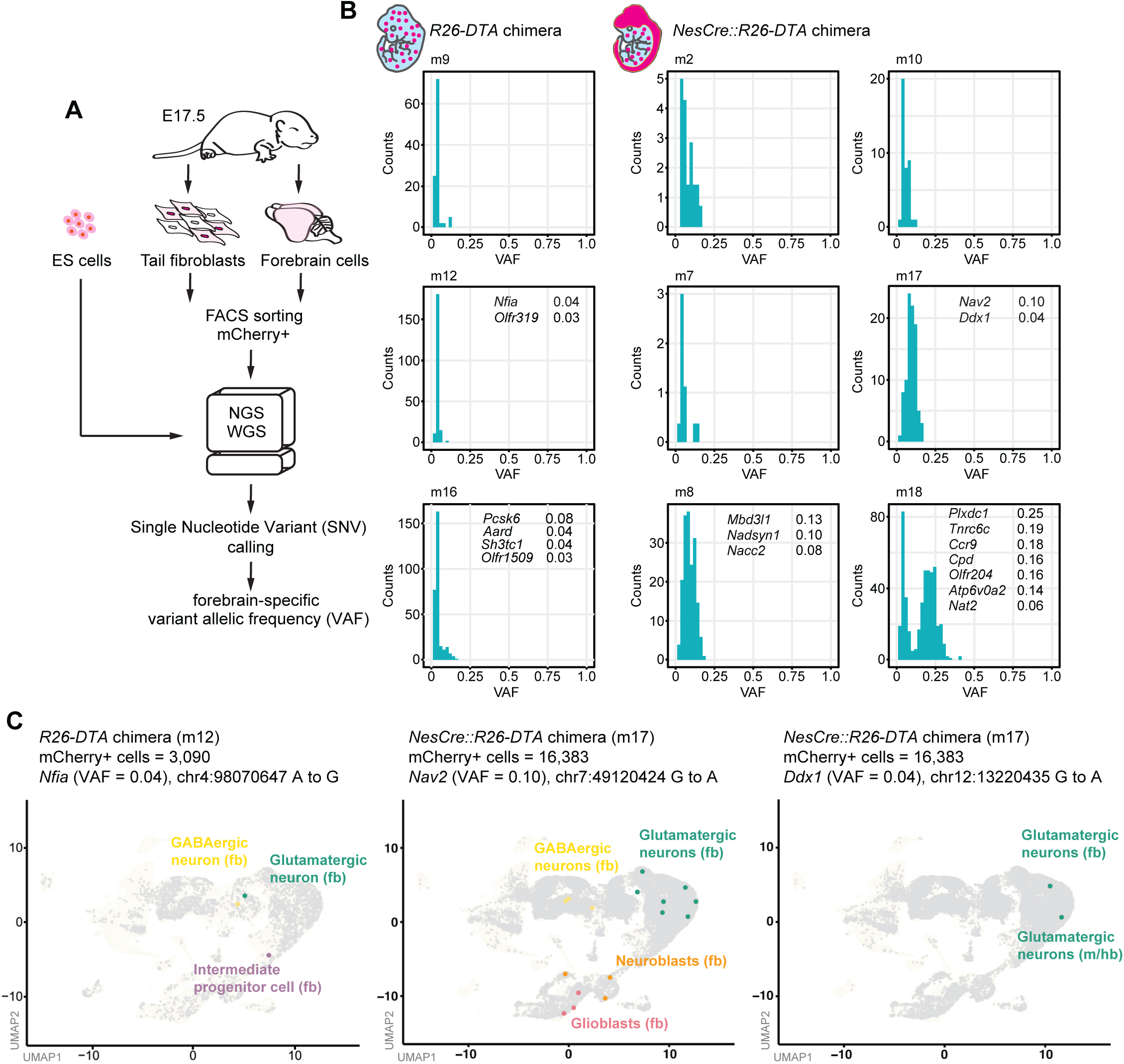
Adaptive NSPC clonal expansion in the *NesCre::R26-DTA* chimeras. **(A)** Schematic of the experimental workflow. At E17.5, tail fibroblasts and forebrain cells were isolated from *R26-DTA* or *NesCre::R26-DTA* chimeras. mCherry+ cells were sorted by FACS and analyzed by whole-genome sequencing (WGS) at a mean of 76x (tail fibroblasts) or 198x (forebrain cells) coverage. Shared allelic variants identified in tail fibroblasts and in ES cell genomes were excluded from analysing forebrain-specific variant frequencies. **(B)** Distribution of variant allele frequencies (VAFs) in mCherry+ forebrain cells from *R26-DTA* (left, n=3) and *NesCre::R26-DTA* (right, n=6) chimeras. Each panel represented an individual mouse (e.g., m09, m12). Exonic and intronic mutations in genes and their frequencies were annotated for select samples. **(C)** The Uniform Manifold Approximation and Projection (UMAP) plots display the distribution of single cells carrying specific clonal SNVs. For each SNV, the VAF determined by WGS, the genomic position, and the substitution type were annotated. Single-cell types carrying intronic SNVs in *Nfia* (left), *Nav2* (middle), and *Ddx1* (right) were indicated. The region of origin of these cells (fb: forebrain; m/hb: mid-/hindbrain), was also noted. The size of the icons represented the expression levels of the indicated genes.

To track clonal evolution after germ-layer separation, we analysed the allele frequencies of variants that were absent in both tail fibroblasts and ES cell genomes. For variants with allele frequencies (VAF) above 2%, we observed a characteristic 1/VAF² distribution, which is typical of subclonal variants^29^ - mutations found in only a subset of the sequenced cells - regardless of the chimera type (**Fig. 2B** at lower frequencies; **Table S3**). These variants represent neutral mutations that accumulated in cells between E6.5 and E17.5. However, in *NesCre::R26-DTA* chimeras only, we identified an unusual cluster of subclonal variants at approximately 10-25% frequency, which was more pronounced in chimera m18 and less so in chimera m7 (**Fig. 2B**, right panels). Since mCherry-positive cells had a diploid genome, clonal mutations (those present in the most recent common ancestor) were expected to appear at a VAF of 50%, reflecting a mutation in one of the two alleles. The presence of these subclones at a VAF of 25% suggested that a dominant neuronal clone made up half of the sequenced cerebral cells, which is extraordinary.

Given that these subclonal clusters of variants were found only in to the *NesCre::R26-DTA* chimeras, they likely represent an early clonal expansion triggered in response to the selectively induced NSPCs death. A complementary multivariate analysis with a variational Binomial mixture model (VIBER^30^) confirmed that four of the six *NesCre::R26-DTA* chimeras (m2, m7, m8 and m18) developed distinct clones (**Fig. S2A**, cluster C2), separated from the neutral low-frequency variants (**Fig. S2A**, cluster C1). To elucidate whether these subclones could be putatively driven by shared mutations, we analysed the nonsynonymous SNVs and indels at the coding regions. We found no known driver somatic mutations shared between *NesCre::R26-DTA* chimeras. In addition, the clonal mutations we found could not explain the expansion phenotype, suggesting they are passenger mutations **(Fig. 2B)**. For the remaining chimeras, namely m10 and m17, VIBER did not recognize a subclone, likely due to the population being a mix of clonal lineages.

In addition to WGS, we also performed scRNA-seq on an *R26-DTA* chimera (m12) and an *NesCre::R26-DTA* chimera (m17). We analysed the somatic SNV in single cell transcripts in order to validate if subclonal SNVs propagated into the neuronal cell. Despite the limitations in sampling and sequencing coverage of the scRNA-seq assay, we found five clonal SNVs across neuronal lineages in the *R26-DTA* and two clonal SNVs in the *NesCre::R26-DTA* chimeras (**Fig. 2C, Table S4**). In the forebrain of the *NesCre::R26-DTA* chimera, a subclone containing an intronic SNV at the *Nav2* gene locus was found in glioblasts, neuroblasts, and GABAergic as well as glutamatergic neurons. The SNV at the first exon of *Ddx1* was found in glutamatergic neuronal populations in both forebrain and mid-/hindbrain, indicating this clone was established at the beginning of neurogenesis (**Fig. 2C**). The number of RG and IP cells expressing *Nfia*, *Nav2,* and *Ddx1* were low (**Fig. S2B**), implying under-sampling of the founders carrying these SNVs.

To eliminate the possibility that the observations resulted from intrinsic nucleotide variations present in the ES cells’ genome, we performed reversed complementation experiments. Instead of injecting wild-type ES cells into the *NesCre::R26-DTA* blastocysts, we injected *NesCre::R26-DTA* ES cells into DsRed-expressing wild-type blastocysts (**Fig. S2C**). In the reversed complementation setting, the surviving cells were derived from the wild-type blastocysts and they globally expressed DsRed (see Material and Methods for more details). All three embryonic brains that were deeply whole-genome sequenced (mean coverage of 180x for the forebrains and 31x for the tail fibroblasts; **Table S2**) displayed an overrepresentation of variants in the DsRed-positive wild-type neuronal cells at E17.5 (**Fig. S2D**). Hence, we concluded that the selective clonal expansion that arose during neurogenesis was independent of pre-existing nucleotide variants in the ES cells’ genome.

In summary, upon induced cell death, the surviving RG cells compete with one another through active clonal expansion, independent of single nucleotide variation or indels present in genes or regulatory elements driving the process.

### Modelling cell cycle dynamics predicts neuronal compensation from few RG clones

The observation that half of the brain originated from a single dominant clone was remarkable. This finding prompted us to develop a mathematical framework to predict clonal expansion dynamics throughout the embryonic neurogenesis period *in silico*.

Our model simulated the normal proliferation dynamics of NSPC populations, comprising of the neuroepithelial cells (NECs), RG cells, IP cells, and the neurons they produce. This model simulated three differentiation steps of the developing mouse brain. First, at E12.5, the model assumed that NECs maintain a constant population size while gradually differentiating into RGs. Second, from E12.5 to E18.5, RGs either self-renew or differentiate into IPs. Third, from E12.5 to E18.5, IPs either self-renew or produce two neurons. To simulate the transition from proliferation to differentiation, the model decreased the self-renewal rate of RGs with the total cell count, eventually approaching zero as neurogenesis progresses. This regulation, based on the total number of progenitors and neurons, ensured that brain development reaches its expected final cell counts **(Fig. 3A)**.

**Figure 3.**
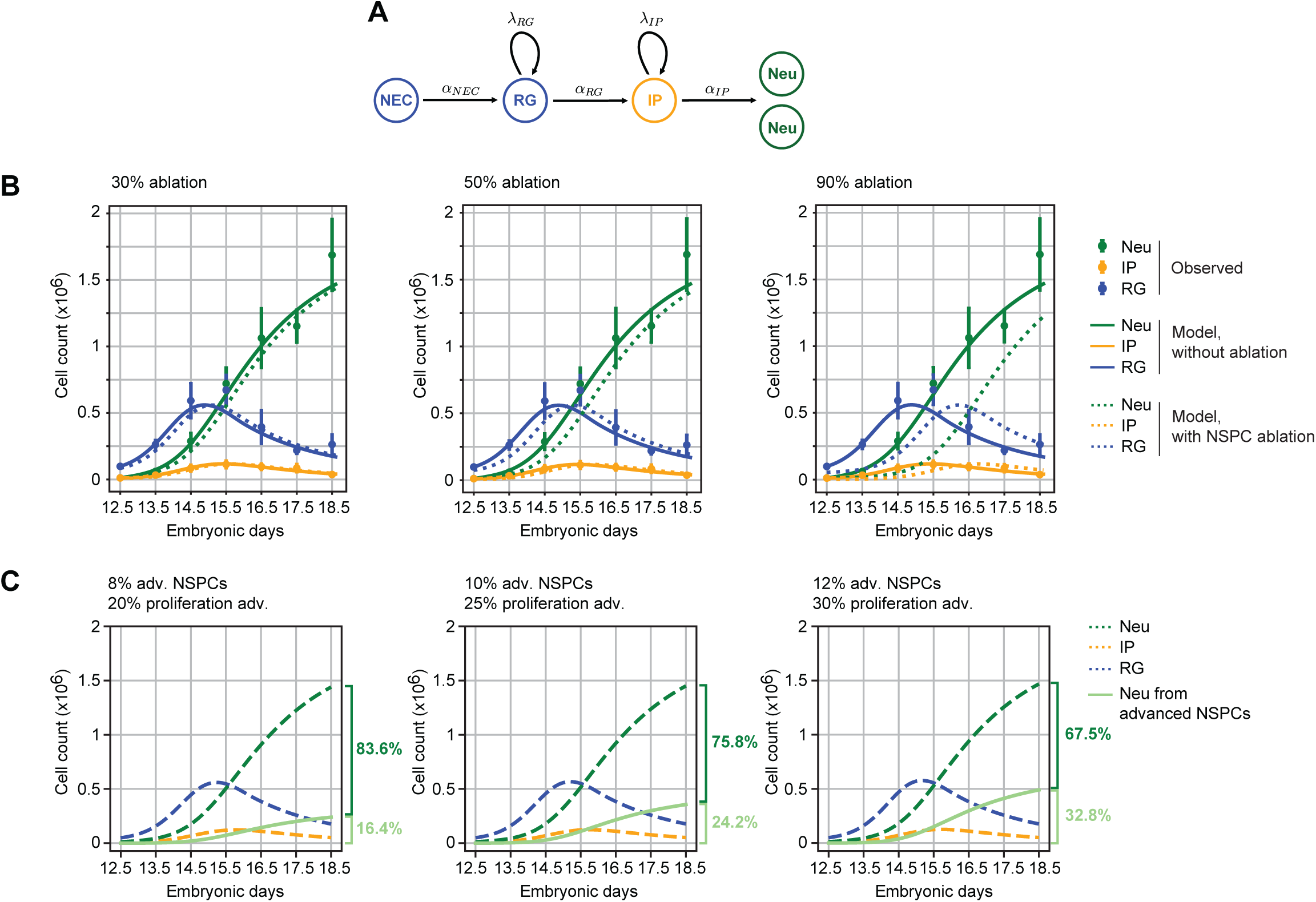
Modelling adaptive clonal expansion. **(A)** Schematic representation of the framework used for modelling NSPC proliferative dynamics and neuron production. NEC: neuroepithelial cells, RG: radial glial cells, IP: intermediate progenitors, Neu: Neurons. 𝛂: differentiation rate. 𝝀: proliferation rate. **(B)** Simulation of progenitor and neuron dynamics under varying levels of RG and IP ablation (30%, 50%, and 90%). Colored dots with the standard deviation represented observed cell counts by Jungas et al.^31^ , while lines represented modelled cell counts under non-ablation (solid) and RG-ablation conditions (dashed). **(C)** Simulation of clonal expansion dynamics with advanced (adv) clones starting at 8% (left) or 12% (right) of the RG. Dashed lines represented cell counts under 50% ablation adjusted with clonal proliferation advantage, as shown in the middle plot in panel B. Light green solid lines represented cell counts derived from clones with proliferation advantages (20%, 25%, or 30%). The percentage of neurons derived from the advanced NSPCs (light green) and the rest of the NSPCs (dark green) at E18.5 was indicated for each simulation.

The model was calibrated using observed cell count data^31^ from the embryonic mouse cortex (**Fig. 3B**). It accurately captured NSPC dynamics and neuron output (**Fig. 3B**, solid lines). Using these rates, we then simulated cell count dynamics under varying levels of NSPCs ablation, ranging from mild (30% and 50%) to severe (90%). Under mild ablation conditions (**Fig. 3B**, left and central panel), the model predicted a half-day delay in RG and IP recovery, accompanied by an accelerated proliferation rate of the surviving progenitors. Despite this delay, neuron numbers closely matched observed values by E18.5. Even under severe ablation (**Fig. 3B**, right panel), the model predicted a delayed recovery of RG and IP cell populations and suggested neuron production could still reach 83.3% of normal levels by E18.5. These predictions indicate that a nearly complete mouse brain could still develop starting from just 10% of the original RG cell population at the onset of neurogenesis.

Next, we simulated how initial clone size and proliferation advantage affect cell competition during neurogenesis (**Fig. 3C**). We started with a clonal size comprising either 8%, 10%, or 12% of the progenitor pool at E12.5 (**Fig. 3C**), adjusting for compensatory reductions in the proliferation rates of non-clonal cells to maintain overall progenitor numbers. Our model predicted that the proliferation advantage plays a critical role in clonal contribution. Clones with higher proliferation rates expanded exponentially, hence allowing even smaller initial clones to achieve dominance. In contrast, lower proliferation advantages slowed clonal expansion, resulting in a more balanced distribution between clonal and non-clonal progenitors.

To experimentally evaluate the cell cycle length, we performed EdU-BrdU pulse-chase labelling *in vivo* at E15.5^32,33^. We chose E15.5 as time point because it exhibited the greatest differences in NSPC counts when ablation was simulated *in silico*. In the developing mouse brains at this stage, the majority proliferating cells were IPs. We found that the cell cycle length of IPs was reduced from 10 to 8 hours in the *NesCre::R26-DTA* E15.5 brains, primarily due to a shortening of the G1/G2/M phases (**Fig. S3A-C**). Considering that IPs in *NesCre::R26-DTA* embryonic brains proliferated two hours faster at E15.5, we predicted that the overrepresented sub-neuronal clones originated from 8-12% of RGs with a 20-30% growth advantage.

These simulations demonstrate that even small clones, when endowed with a proliferation advantage, can disproportionately grow to dominate the progenitor pool, highlighting the system’s capacity for compensating neurogenesis while maintaining the overall cell population dynamics.

### C>A and C>T substitutions characterized the *NesCre::R26-DTA* chimera’s genomes

To investigate the underlying processes promoting clonal expansion, we used single-base substitution (SBS) analysis, which examines mutations where one nucleotide is replaced by another in the DNA sequence. SBS patterns predict distinct mutational signatures, providing insights into the processes contributing to genome alterations in both normal and cancer tissues^34^. In mCherry-positive neuronal cells isolated from the *R26-DTA* chimeric embryos, most of the SNVs are C>T or T>C substitutions, typically associated with aging and cell cycles^34^ (**Figs. 4A, B**). In contrast, C>A substitutions were significantly enriched at the mCherry-positive neuronal genomes in the *NesCre::R26-DTA* chimeras. C>A substitutions are strongly linked to oxidative DNA damage^34^. In the reversed complemented chimeras, most of the SNVs were C>T substitutions, with fewer C>A substitutions (**Figs. S4A, B**).

**Figure 4.**
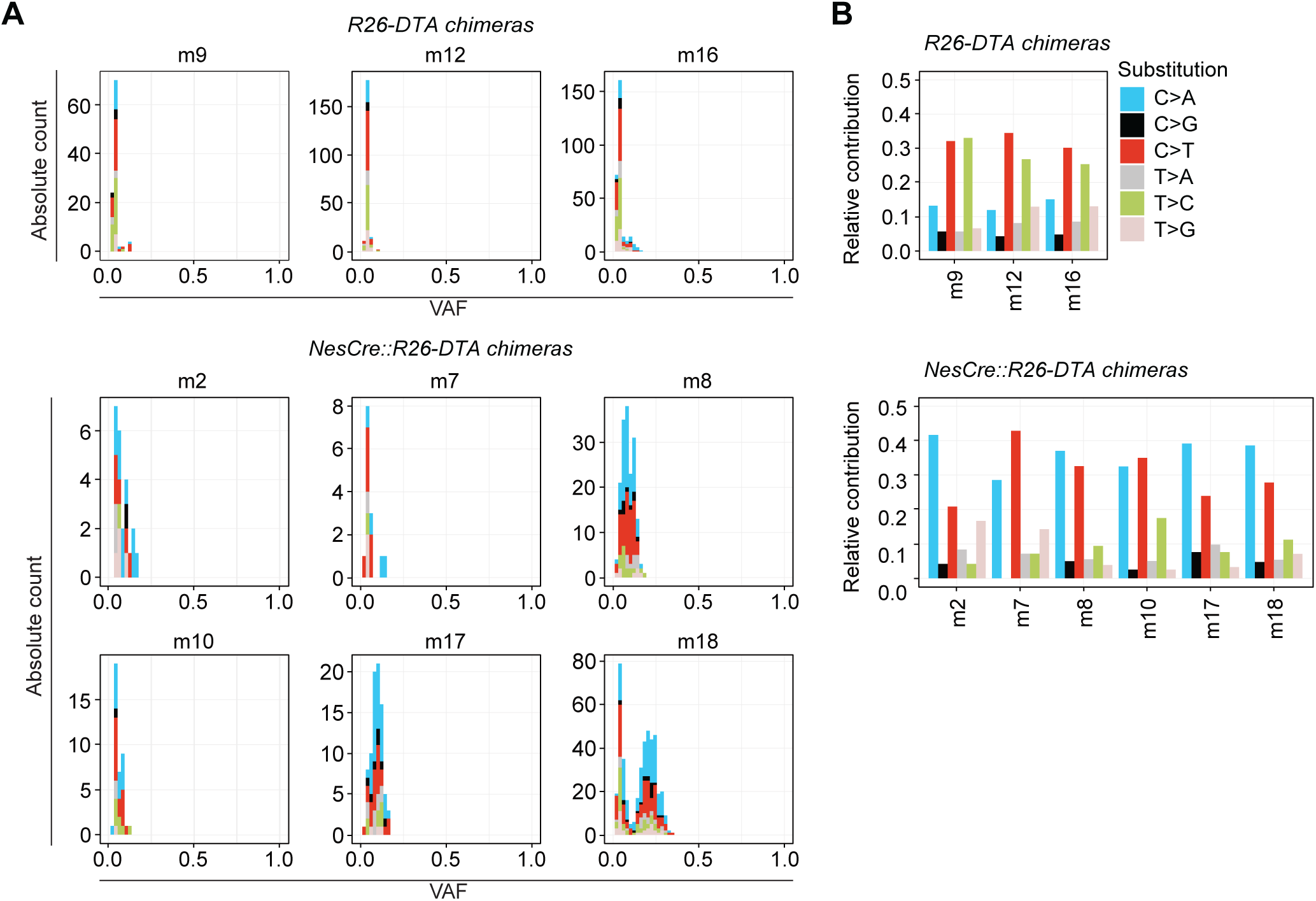
Mutational signatures in *R26-DTA* and *NesCre::R26-DTA* chimeras revealed distinct patterns of single base substitutions. **(A)** Variant allele frequency (VAF) distributions for each substitution type (represented by different colors and stacked) in mCherry+ forebrain cells of *R26-DTA* chimeras (top: m09, m12, m16) and *NesCre::R26-DTA* chimeras (bottom: m02, m07, m08, m10, m17, m18). The *NesCre::R26-DTA* chimeras displayed broader VAF distributions. **(B)** Relative contribution of each substitution type across individual *R26-DTA* (top) and *NesCre::R26-DTA* (bottom) chimeras. *NesCre::R26-DTA* samples showed an increased proportion of C>T and T>A mutations compared to controls.

In summary, SBS analysis indicated that most NSPCs underwent more cell cycles (as evidenced by the C>T substitutions) and, in some cases, engaged biological processes that generate oxidative stress, which possibly promoted clonal expansion.

### Adaptive clonal expansion slowed neural stem/progenitor cell fate transition

Given that a few subclones formed an entire brain within just five embryonic days, we wondered whether the neuronal cells in the brains of *NesCre::R26-DTA* chimeras had a biological age equivalent to those in *R26-DTA* chimeras. To address this, we analysed the frequency of various cell types in the chimeras. In the mCherry-positive cell populations of *NesCre::R26-DTA* chimeric embryos, we observed a reduced proportion of Cajal-Retzius cells, ependymal cells, RG cells, glioblasts, and neuroblasts in the brain at E17.5 (**Fig. 5A**). Specifically, the combination of RG cells, glioblasts, and neuroblasts - representing the NSPCs at this stage - decreased from 16% to 14.5%. We reasoned that this was because of the NSPCs actively producing neuronal progenies through asymmetric proliferation and differentiation, thereby consuming themselves. Indeed, 78% of cells in the *NesCre::R26-DTA* embryonic brains were neurons, which was six percent higher than in *R26-DTA* embryos. The other neuronal cell types did not show significant differences between genotypes (**Fig. S5A**).

**Figure 5.**
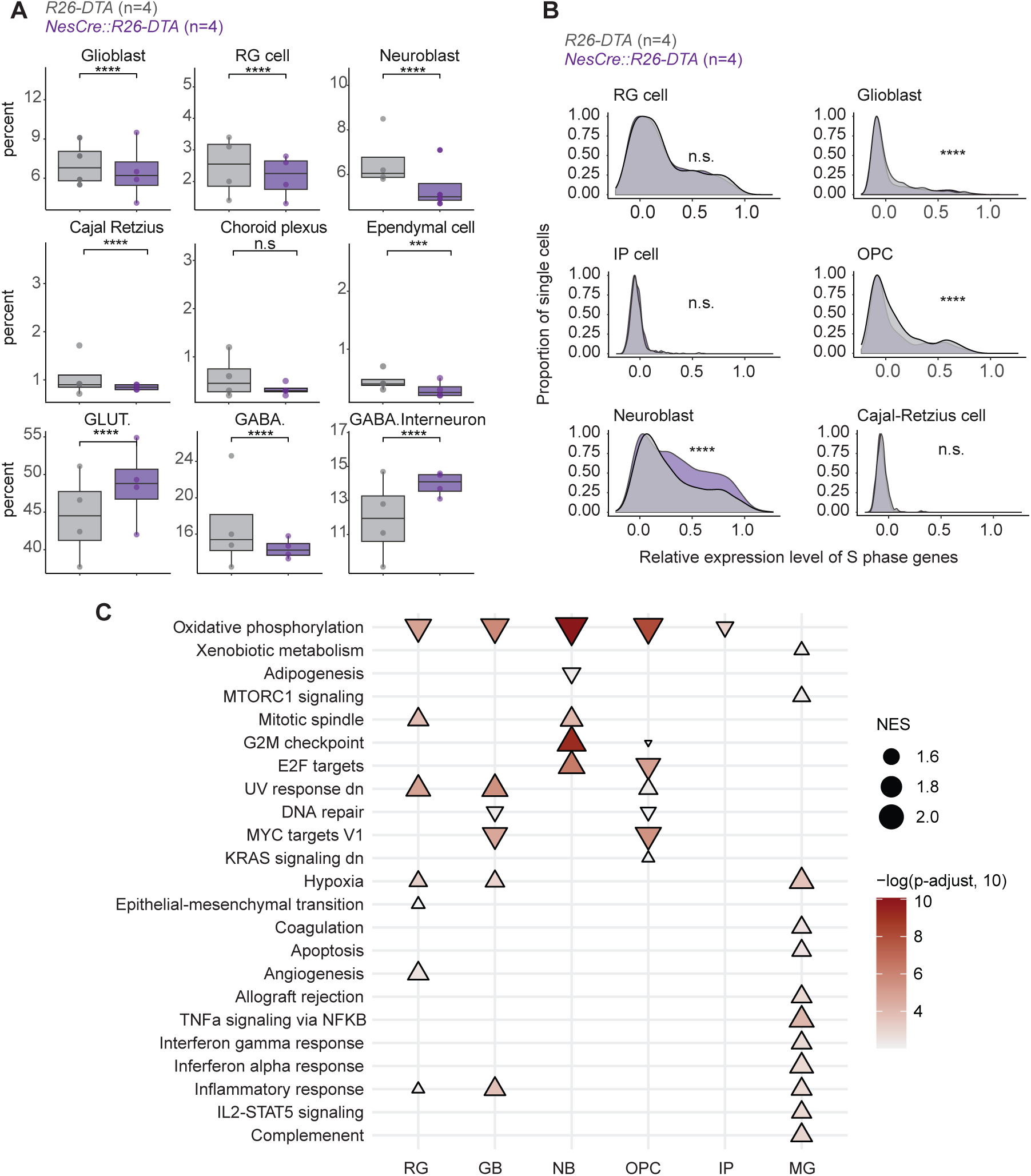
Adaptive clonal expansion delayed NSPC cell fate transition and altered cell composition in *NesCre::R26-DTA* chimeras. **(A)** Proportional analysis of brain cell types in E17.5 *NesCre::R26-DTA* and *R26-DTA* chimeras. Four biological repeats for each condition were analysed. Statistical significance was determined by a chi-square test. *** p ≦ 0.001, **** p ≦ 0.0001. **(B)** S phase score in NSPCs. All mCherry+ single cells were considered for this analysis. The x-axis depicts the activity of S phase genes, determined by CellCycleScore of the S phase, and the y-axis indicates the proportion of single cells in each cell type. Non-cycling cells showed a peak centered at 0 on the x-axis (e.g. Cajal Retzius cells). Statistical significance was determined by a Kolmogorov–Smirnov test. **** p ≦ 0.0001. n.s.: not significant. **(C)** Gene set enrichment analysis was conducted to compare *NesCre::R26-DTA* and *R26-DTA* chimeras for each cell type based on the scRNA-seq dataset. The direction of the triangle indicates whether a pathway is up- (▴) or downregulated (▾) in the *NesCre::R26- DTA* compared to the *R26-DTA* chimeras. The size of the triangle depicted the normalized enriched score (NES). The color indicated the adjusted p-value, shown on a scale of -log 10. RG: radial glial cells, GB: glioblasts. NB: neuroblasts. OPC: oligodendrocyte precursor cells. IP: intermediate progenitors. MG: microglia cells.

In addition, the transition of cell fate can be inferred from the proportion of each NSPC type (highlighted in blue in **Fig. S5B**) that remains in the S phase. By E17.5, most RG cells had exited the cell cycle in both *NesCre::R26-DTA* and *R26-DTA* brains (**Fig. 5B**). However, we identified a significantly higher proportion of neuroblasts that were still in the S phase within the *NesCre::R26-DTA* brains, accompanied by a reduction in S phase activity in oligodendrocyte precursor cells and glioblasts (**Fig. 5B**). Since oligodendrogenesis and gliogenesis typically occur after neuroblast-dependent neurogenesis^35^, we speculated that embryonic neurogenesis was delayed.

Furthermore, we found the expression levels of genes associated with oxidative phosphorylation were reduced in the *NesCre::R26-DTA* NSPCs – namely, RG cells, IPs, glioblasts, neuroblasts, and oligodendrocyte precursor cells (**Fig. 5C, Fig. S5C-E**). As NSPCs shift from aerobic to oxidative metabolism during cell fate transitions^36^, low oxidative phosphorylation suggested a delay in the cell fate transition process. Lastly, the expression levels of genes involved in the G2/M checkpoint pathway in the *NesCre::R26-DTA* NSPCs were higher than in the *R26-DTA* controls. This indicated that the fast-cycling NSPCs produce more DNA damage during the S phase, further suggesting that cells experienced replication stress. Conversely, microglia cells, lymphoid-origin cells migrating to the mouse brain at E9.5^37^, did not show alterations in oxidative phosphorylation or in the G2/M checkpoint pathways, suggesting the overall phenotype was limited to the neuronal lineage (**Fig. 5C**).

In summary, although NSPCs reduced cell cycle length to meet the demands of brain development, they experienced a delayed cell fate transition at the end of embryonic neurogenesis. Adaptive clonal expansion also consumed the pool of NSPCs, leaving significantly fewer NSPC reserves for life.

### Limitations of the study

Although *Nestin* mRNA is detectable in the central nervous system as early as E7.5^38^, the transgenic rat *Nestin* promoter and enhancer used to drive Cre expression in this study only become active from E12.5 onward^22^. Furthermore, the neuron output prediction model relies on existing neuronal cell count data, which is unavailable between E10.5 and E12.5. As a result, the model is constrained to address clonal competition starting from E12.5 and not earlier. Consequently, our analysis did not incorporate clonal expansion between the germ layer branching point and the onset of neurogenesis.

## Discussion

In this study, we showed that the reduction in the NSPC population created a stochastic developmental bottleneck, which allowed NSPCs to unleash their proliferation potential. This scenario promoted adaptive expansion of a small subset of NSPCs, ultimately restoring the neuronal cell types despite significant brain depletion (**Fig. 1**). Variant allele frequency analysis provided evidence of selective clonal expansion, occurring independently of known driver genetic mutations (**Fig. 2**). We hypothesize that an epigenomic or metabolomic adaptation enabled NSPCs to compete and expand clonally when neighbouring cells are lost. Our findings also showed that clonal dominance during neurogenesis depends on both initial clone size and proliferation advantage, highlighting the neurogenic system’s adaptability to balance clonal expansion with overall progenitor pool stability (**Fig. 3**). This expansion is associated to biological processes that generate oxidative DNA damage, a by- product of accelerated proliferation (**Fig. 4**). We further speculate that these clonal expansions occurred between E10.2 and E12.5, a timeframe for which we lacked accurate measurement data for fitting. While clonal expansion in the human brain has been reported^12,13^, the underlying reasons and the potential health consequences remain unclear. Our findings suggest that this driver-independent expansion may be associated with a developmental crisis affecting the NSPC pool size and may have impact on brain diseases.

While neuron numbers were restored through rapid NSPC expansion (**Figs. 1,3**), this process came at a cost. We observed delayed cell fate transitions and a significant depletion of NSPC reserves by the end of embryogenesis, implying consequences for postnatal brain development and long-term neurogenesis (**Fig. 5**). This observation aligns with earlier studies in adult mice, which showed that certain neural stem cells, typically dormant, become activated under stress but fail to re-enter the cell cycle after a second wave of loss, suggesting an irreversible dormancy state^39^. Our findings suggest that adaptive clonal expansion may consume these stem cell reservoirs, posing risks for impaired neurogenesis later in life.

Contrary to the lateral inhibition model, which proposes that Notch1-Dll1 interactions promote self-renewal and suppress asymmetric neurogenesis^40^, we found that losing neighboring cells facilitated RG cells to restore the progenitor pool. This shift likely occurs due to reduced competition for Notch1 ligands, enabling the remaining RG cells to re-enter active cycling. Alternative pathways, such as FGF2 signalling, hypoxia-induced cell cycle acceleration, or soluble Notch1 ligands, may complement this framework. Furthermore, progenitor cell pool reduction-driven clonal expansion can also be observed in real-world scenarios, such as hematopoietic cell transplantation. In this context, clonal diversity declines after allogeneic transplants due to selective pressures or loss of a suppressive niche^41^ . Unlike the foreign environment hematopoietic stem cell encountered in the recipients, adaptive clonal expansion in the brain reflects a native stem cell response to developmental stress.

## Material and Methods

### Mice

All mice were bred in-house at the DKFZ Center for Preclinical Research and maintained under specific pathogen and opportunistic-free conditions on a 12-hour light-dark cycle in a temperature-controlled environment with *ad libitum* access to food and water. All procedures adhered to DKFZ guidelines and were approved by the Regierungspräsidium Karlsruhe (G154/20 and G241/20). To generate brain-ablated embryos, NesCre male mice (B6.Cg-Tg(Nes-cre)1Kln/J, Jackson Laboratory) were crossed with R26-DTA female mice (B6.129P2-Gt(ROSA)26Sortm1(DTA)Lky/J, Jackson Laboratory), producing *NesCre::R26-DTA* and *R26-DTA* embryos. Mating females were checked for vaginal plugs (VP) the following morning, and pregnant females were euthanized via cervical dislocation 12- or 15- days post-mating to collect E12.5 and E15.5 embryos. SW nursing females and vasectomized SW males were bred in-house at DKFZ for blastocyst complementation. DsRed (B6.Cg-Tg(CAG-DsRed*MST)1Nagy/J) mice were used for reversed complementation experiments following the above mentioned procedures.

### Blastocysts Complementation

Blastocyst injection and implantation followed procedures described in ^42^, with minor modifications. In brief, 6- to 12-week-old R26-DTA females were used as blastocyst donors. They were superovulated by intraperitoneal injection of 5 I.U. pregnant mare serum gonadotropin (PMSG), followed by 5 I.U. human chorionic gonadotropin (hCG) 46-48h later. *R26-DTA* homozygous females and *Nestin* hemizygous male animals were set in the same cage at the earliest one hour after the last hormone administration. The next morning, the mated females were checked for VP. 3.5 days post-mating, which was equal to day 2.5 for morulae collection, VP-positive females were killed by cervical dislocation. The oviducts and part of the uterus were removed, and the blastocysts, or morulae, were isolated by flushing the oviduct/uterus. On day 3.5, around 15 mCherry-positive mouse ES cells from the cell line JM8/A4 were injected into the blastocyst cavity. These blastocysts were then implanted bilaterally into the uteri of a pseudopregnant uteri of a 2.5-day pseudopregnant Swiss Webster (SW) female mouse. At E15.5 or E17.5, the pregnant SW females were killed by cervical dislocation, and the chimeric embryos were harvested.

For the reversed complementation experiments, 6- to 12-week-old *DsRed::R26-DTA* homozygous females were used as donors. These females were superovulated and mated with *Nestin* hemizygous males. Blastocysts were isolated, consisting of either *DsRed::R26- DTA* hemizygous or *DsRed::NesCre::R26-DTA* hemizygous genotypes. *NesCre::R26-DTA* mouse ES cells were then injected into all blastocysts. All *DsRed:: NesCre::R26-DTA* chimeras exhibited complete ablation phenotypes, leaving no brains for analysis. Deep WGS was performed on DsRed-positive cells derived from *DsRed::R26-DTA* chimeras.

### Measurement of Cell Cycle Length Using BrdU/EdU Pulse-Chase *in Vivo*

Pregnant female mice carrying embryos at E15.5 were used for the *in vivo* pulse-chase experiments. At embryonic day E15.5, the pregnant female mice received an intraperitoneal injection of 5-ethynyl-2′-deoxyuridine (EdU) at a dose of 50 mg/kg body weight followed by 5- bromo-2′-deoxyuridine (BrdU) injection at 100 mg/kg body weight after a chase period of one hour. EdU and BrdU were both dissolved in sterile PBS and administered separately to label S-phase cells at two distinct time points. 30 minutes after BrdU injection, the mice were euthanized, the embryos were isolated and fixed in 10% (w/v) neutral buffered formalin solution for at least 48 hrs at RT. The embryos were then FFPE processed and cut as described in the embryo preparation for histology and Immunofluorescence section below.

TBR2-, BrdU-, or EdU-positive cells were determined by immunofluorescence staining. The number of TBR2+ cells that were either positive or negative for BrdU or EdU, respectively, as well as triple positive cells, were determined by using the neural network-based cell segmentation software CellPose2^43^ in combination with a custom ImageJ Macro workflow for analysing marker expression and their colocalization. In Cellpose, we trained custom models and adapted the flow and the cell probability threshold for each marker of interest, based on Cellpose’s integrated nuclei model^44^. To calculate cell cycle length and S phase length, we adapted the published approach^32^, using the following formulas:

- S-phase-length = BrdU+/BrdU-EdU+
- cell cycle length = S-Phase-length/(BrdU+/all).

### Cell Sorting

To enrich mCherry-positive neuronal cells and tail fibroblasts, pseudopregnant SW females were euthanized by cervical dislocation at E17.5. After dissection, embryos were immediately placed in PBS 1X solution on ice. The forebrain was microdissected, the meninges removed, and the olfactory bulb was dissected out. For single-cell RNA sequencing experiments, the forebrain and mid hindbrain from each embryo were micro-dissected and used as one replicate. For WGS experiments, each forebrain isolated from one embryo was used as one replicate and tail embryonic fibroblasts from the corresponding embryo were isolated and used as a control. Brain tissue was digested using the papain dissociation kit (Worthington Biochemical Corporation, LK003150), and tail embryonic fibroblasts were obtained by digesting the embryonic tails using a digesting solution consisting of pronase and collagenase D, following the published protocol^45^ .

#### Mouse Embryo Preparation for Histology and Immunofluorescence Sample Collection

Mouse embryos were collected at E12.5 and E15.5 after timed mating. Pregnant females were euthanized by cervical dislocation following institutional animal care guidelines. Embryos were dissected in cold phosphate-buffered saline (PBS) and immediately processed for fixation.

#### Fixation

Embryos were fixed in 10% (w/v) neutral buffered formalin solution for at least 24 hours at room temperature. After fixation, embryos were then processed for dehydration on a STP 120 processing machine (Thermo Fisher) and embedded in paraffin using a Histo Star embedding station (Thermo Fisher) for histological sectioning. Sections (thickness: 6 µm) were cut using a RM2255 microtome (Leica), mounted on Super-Frost Plus glass slides, and dried overnight at 37°C.

#### H&E Staining

Paraffin sections were deparaffinated and rehydrated as follows: 2x 10 min in xylene, 2x 10 min in 100% ethanol, 2x 5 min in 95% ethanol, 2x 5 min in 70% ethanol and 1x 5 min in ddH2O. The staining was performed with the deparaffinated and rehydrated sections by the following steps: 30 sec in hematoxylin solution, 20 sec rinsing with tap water, 10 sec in 0.1% hydrochloric acid, 6 min rinsing with warm tap water, 30 sec in eosin solution, 1 min rinsing with flowing tap water. After the staining, dehydration of the slides was performed as follows: 1x 3 min in 70% ethanol, 1x 3 min in 85% ethanol, 2x 5 min in 100% ethanol and 2x 5 min in xylene. Using Eukitt mounting medium (ORSAtec GmbH #202200401), the dehydrated and stained slides were mounted using glass coverslips. Imaging was performed afterwards by using an Axioscan 7 slide scanner (Zeiss).

#### Immunofluorescence Staining and imaging acquisition

Sections were deparaffinized and rehydrated as described in the H&E staining section, then washed three times in 1X PBS solution. Antigen retrieval was performed by boiling the sections at 80°C for 10 minutes in a solution of citrate buffer (BioGenex #HK088-9K; 25 mL of 10X citrate buffer diluted with 225 mL of ddH₂O).The sections were blocked in 10% normal donkey serum in PBS-T (with 0.2% Triton X-100) for 1 hour, RT. Click-IT reaction was performed using the Click-iT™ EdU Cell Proliferation Kit for Imaging, Alexa Fluor™ 647 dye (ThermoFisher Scientific cat. NO. C10340) according to the manufacture instruction to detect EdU. Sections were then incubated with primary antibody against BrdU (rat, Abcam ab6326) at 4°C overnight. After washing with PBS, sections were incubated with the fluorescently conjugated secondary antibody anti-rat Alexa Fluor® 750 (goat, Abcam ab175751) for 1 hour, RT. The next day, the same sections were incubated with primary antibody against PAX6 (rabbit, Biolegend, cat #901301) or fluorophore-conjugated TBR2 (Alexa488, rat, eBioscience 53-4875-82) primary antibody, and a fluorophore-conjugated secondary antibody (Alexa 555, goat-anti-rabbit, ThermoFisher A32732) was used to label primary antibody against PAX6. After washing, nuclei were counterstained with DAPI. Slides were mounted with ProLong Gold Antifade Mounting Reagent using glass coverslips. Immunofluorescent images were obtained using THUNDER Imager 3D Live Cell (Leica), using an HC PL APO 20x/0.80 lens. All samples were blinded before the acquisition of the data.

#### Whole-genome sequencing Sample Preparation

Neuronal cells were isolated from the E17.5 mouse embryonic brain following the procedure described in the Cell sorting section. For control comparisons, tail fibroblasts and mouse ES cells were harvested following established protocols. mCherry-positive cells were sorted by using Aria, Fusion, and Symphony S6 SE, and genomic DNA was extracted from all samples using the PureLink™ Genomic DNA Mini Kit (Invitrogen cat. N. K1820-00) according to the manufacturer’s instructions. DNA quality and concentration were assessed using a NanoDrop spectrophotometer and Qubit dsDNA Assay Kit, respectively.

#### Whole-Genome Sequencing

Whole-genome sequencing libraries were prepared using the Illumina DNA Prep, (M) Tagmentation kit, and the libraries were mixed to create a multiplex. Pair-end sequencing was performed using the NovaSeq 6000 S4 V1.5 or the NextSeq 500 Mid-Output sequencer to achieve a depth of approximately 50X for ES cells, 30X for tail fibroblasts, and 150X for neuronal cells (see **Table S2** for sequencing coverage details). Sequencing reads were 150bp in length, and a minimum of 52 million reads, 140 and 255 million reads per sample were generated for tail, forebrain and ES cell line, respectively.

#### Germline and Somatic SNV Calling

BAM files were generated from the raw sequencing data using BWA (version 0.7.15) to align reads to the reference genome GRCm38mm10_PhiX with the help of the DKFZ ODCF team. Somatic single nucleotide variants (SNVs) and small insertions/deletions (indels) in the forebrain samples were identified using Strelka (v2.9.244) and Mutect2 (GATK v4.2.0.043), with matched tail fibroblast samples and ES cell sequencing, both serving as germline controls. Variants located in repeat and simple repeat regions were filtered out using Bedtools Intersect (v2.24.045). High-confidence variants were defined as those passing the default filters of both Strelka and Mutect2.

#### Variant Allele Frequency

Variant allele frequencies were calculated for each small variant (SNVs and indels) using GATK. Only variants with a VAF ≥ 0.02 in neuronal cells and absent in both tail fibroblasts and ES cells were included for further analysis. The neuronal-specific SNVs were then used in downstream mutation signature and single base substitution (SBS) analyses.

#### Binomial clustering of VAF

To perform binomial clustering we use VIBER, where we require the minimum cluster size to be 2% of the total number of points.

#### Single Nucleotide Substitution and Mutation Signature

To investigate the mutational processes in neuronal cells, SBS patterns were categorized into the six classes of base substitutions (C>A, C>G, C>T, T>A, T>C, T>G). Mutation signature analysis was performed using the COSMIC database.

Somatic SNVs were mapped to these signatures, and the de novo extraction of mutational signatures and relative contribution of each signature was calculated using SigProfilerExtractor^46^ .

#### Mathematical Modeling of Neural Progenitor Dynamics Model Framework

A system of ordinary differential equations (ODEs) was developed to simulate neural progenitor cell types’ proliferation and differentiation dynamics during early brain development. The model tracks neuroepithelial cells (NECs), radial glial cells (RGs), intermediate progenitors (IPs), and the neurons (Neu) they produce.

This model assumes that NECs self-renew to maintain a constant population size. They then differentiate into RGs at rate 𝛼_NEC_. RGs self-renew at rate 𝜆_RG_ and differentiate into IPs at rate 𝛼_RG_. The RGs’ self-renewal rate, 𝜆_RG_ , decreases over time, depending on the total cell count 𝑛_total_. This aims at simulating the transition from proliferation to differentiation during neurogenesis as the cell population in the neocortex approaches a carrying capacity 𝐶𝐶. Lastly, IPs self-renew at rate 𝜆_IP_and divide to produce two neurons at rate 𝛼_IP_. The system of ODEs is given by:

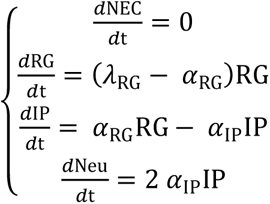

where

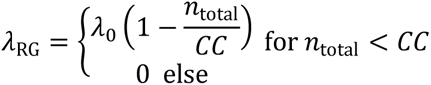

#### Model Calibration and Validation

The model was calibrated using quantitative flow cytometry data^31^ , which provided absolute counts of progenitor cell types from E12.5 to E18.5. The model predicts 𝛼_NEC_ = 𝜆_IP_ = 0, i.e. NECs do not proliferate into RGs after E12.5, and IPs do not self-renew. These predictions were validated by comparing simulated results to observed data across embryonic time points. The model accurately predicts progenitor population dynamics under non-ablation conditions.

#### Ablation and Clonal Dynamic Simulations

The model applied a 30%, 50%, or 90% reduction in NEC, RG, and IP populations at E12.5 to simulate ablation. To simulate clonal dynamics, we took the proliferation and differentiation rates from the model and ran a Gillespie simulation. We assumed an initial clone consisting of between 8%, 10%, and 12% of RG cells at E12.5, proliferating between 20% and 30% faster than the remaining cells, determined by a selective advantage. To keep the total cell count constant, the proliferation rate of the non-clonal cells was adjusted.

#### Single-Cell RNA-sequencing Sample Preparation

The entire embryonic mouse brain was harvested at E17.5. The embryonic brains were dissected under a stereomicroscope, dividing into the forebrain (including the thalamus and the neocortex) and the mid-/hindbrain (including medulla oblongata, cerebellum, and midbrain). The dissected brain parts were dissociated into a single-cell suspension using the papain dissociation kit (Worthington Biochemical Corporation, LK003150) according to established protocols. Cells were filtered through a 70 µm cell strainer to remove debris, and viability was assessed using trypan blue staining to ensure a minimum viability of 90%. Cells were counted, and the concentration was adjusted to 500,000 cells/mL for downstream applications.

#### Library Preparation

Single-cell RNA sequencing libraries were prepared using the 10x Genomics Chromium platform (Single Cell 5’ v2 kit) following the manufacturer’s protocol at the single-cell Open lab at the DKFZ. Briefly, 20,000 of single cells were loaded into the Chromium Controller to generate single-cell gel bead-in-emulsions (GEMs). Reverse transcription was carried out within the GEMs to capture mRNA and generate cDNA. In the reverse transcription step, a custom primer was added to the reaction at 10nM concentration to specifically enrich the mCherry transcript. Libraries were constructed through subsequent amplification, fragmentation, and attachment of Illumina sequencing adapters.

#### Sequencing

The prepared libraries were sequenced on an Illumina Novaseq 6000 S2 to a depth between 50,000 and 60,000 reads per cell. Sequencing was performed in paired-end mode with 28 cycles for Read 1 and 90 cycles for Read 2 to obtain high-quality reads for downstream analyses. The QC plots of each single-cell sequencing sample can be found at the Github repository (https://github.com/orgs/brainbreaks/repositories/NPC-Clone-Plasticity)

#### Data pre-processing and clustering

The raw sequencing data were processed using the CellRanger pipeline (v7.0.0, 10x Genomics), aligning reads to a custom genome reference built using the mkref function in CellRanger. This reference was based on the mouse genome and transcriptome (mm10 from 10X Genomics), with additional sequences for mCherry, DTA, and Cre (included in the GTF and FASTA files). Default parameters were used to generate gene-by-cell count matrices.

Raw count matrices were individually processed to remove doublet cells using Scrublet^47^ . The Seurat package^48^ (v4.3.0) was employed to generate sparse count matrices for downstream analyses. Cells were filtered to retain those with higher quality, defined by the detection of at least 500 genes and a mitochondrial gene expression percentage of ≤5%. Data normalization followed the LogNormalize function in Seurat, using a scaling factor of 10,000. Additionally, 3000 highly variable genes were identified, and the effects of number of detected features (nFeature) RNA and mitochondrial gene expression percentage were regressed out for subsequent analyses.

Principal component analysis (PCA) was performed on the top 3000 highly variable genes, retaining 50 principal components for downstream analyses. To integrate single-cell data across different batches, Harmony^49^ (v1.0.1) was applied. The integrated data were subsequently used to construct a k-nearest neighbors’ graph, followed by clustering with the Leiden algorithm, using a final resolution of four.

#### Cell Type Annotation

To annotate cell types in the single-cell data, we performed high-resolution clustering within each cluster to generate meta-cells for annotation. We evaluated 23 different resolutions, ranging from 1 to 10 in increments of 0.4, and selected the optimal resolution based on the size and nFeatures of the meta-cells. Canonical correlation analysis (CCA)-based label transfer, as implemented in Seurat, was used to transfer annotations from the reference atlas^50^ utilizing the 3,000 highly variable genes identified. The final cluster annotations were determined by evaluating the highest predicted scores, consensus agreement of transferred labels, and the expression patterns of canonical marker genes.

#### Cell Type Proportion Statistical Analyses

We calculated the cell type composition across biological replicates for both the *NesCre::R26-DTA* and *R26-DTA* conditions. To account for the replicated information, we applied a Generalized Linear Model with a binomial distribution followed by an ANOVA using the chi-square test to assess statistically significant differences in cell type proportions between the conditions.

#### S Phase Score Analyses

We assigned cell cycle scores to each cell using the Seurat CellCycleScoring function and assessed the distribution differences between the *NesCre::R26-DTA* and *R26-DTA* conditions within each cell type using the Kolmogorov-Smirnov test.

#### Regulon Analyses

We used SCENIC^51,52^ to infer gene regulatory networks, utilizing the latest motif database (mm10_10kbp_up_10kbp_down from SCENIC) and transcription factors from the cisTarget database for *Mus musculus*. To identify conserved regulons, we implemented the SCENIC algorithm using pySCENIC (v0.12.1) in Python (v3.8.19), running 78 iterations. In each iteration, we performed random subsampling with replacement to obtain 20,000 cells, applying GRNBoost2 with default settings. For consensus regulon identification, we retained regulons with a frequency of occurrence greater than 50% across iterations and used the average weight to perform AUCell scoring for all cells.

#### Pathway Enrichment Analysis

We performed Gene Set Enrichment Analysis (GSEA) using fgsea^53^ (v1.30.0 in R 4.3.0) and the Hallmark gene sets for *Mus musculus* from Msigdbr (v7.5.1) (https://github.com/igordot/msigdbr, also ^54^ ) to compare the *NesCre::R26-DTA* and *R26-DTA* conditions within each cell type. The input for GSEA was derived from the results of a Wilcoxon rank-sum test (wilcoxauc) performed using Presto (v1.0.0.).

#### Regulon Activity and Associated Pathways Analysis

The activity of regulons showing significant differences (p-values ≤0.01) between the *NesCre::R26-DTA* and *R26-DTA* conditions, as identified through a permutation test, was selected for further analysis. To explore pathways associated with each regulon, we used the same Hallmark gene sets as the GSEA analysis. Fisher’s exact test was performed on the Odds Ratio, considering the overlap between regulon and pathway gene sets. As a background, we included gene sets with more than 5% expression in the single-cell population. Pathways and regulons with an Odds Ratio ≥1 and p-values ≤0.01 were considered significantly associated. Regulons with gene set overlaps below the 10th percentile were excluded. To validate that these overlapping gene sets contributed to pathway differences, we applied the Kolmogorov-Smirnov test on module scores (calculated via Seurat’s AddModuleScore function), retaining only those with significant differences (p-values ≤0.01) for the final results.

#### Single-Cell Variant Calling

We utilized CellSNP-lite to perform SNP calling at single-cell resolution using E17.5 10x Cell Ranger outputs. This analysis was restricted to paired animals with whole-genome sequencing (WGS) data (mouse12 and mouse17). SNP calling was conducted in Mode 1a with default settings, except for the --minMAF 0 and -- minCOUNT 1. Additionally, we considered only SNPs that were also identified in the WGS data of paired animals and known to contribute to clonal expansion.

#### Code availability

R-based scripts for analyzing the scRNA-seq data were deposited under the brainbreaks GitHub repository (https://github.com/orgs/brainbreaks/repositories/NPC-Clone-Plasticity). WGS datasets have been uploaded as Bioproject PRJNA1209637 (BioProject and associated SRA metadata are available at https://dataview.ncbi.nlm.nih.gov/object/PRJNA1209637?reviewer=ar8r2fidffnio7pl9rv8ne6m2m) and the R script for analyzing it will be published on Github upon publication. For reviewers: https://tsbnc.dkfz.de/index.php/s/W7RLZYr4WAzWG5w Password: 7S4UrIeRVm.

#### Statistical Power

Statistical power for all the experiments was mentioned in the associated text, figure legends, or methods.

## Acknowledgments

This work is supported by the Helmholtz Young Investigator grant and an ERC starting grant BrainBreaks to P-C W.. G.D.M. received a Conference Travel Program from DAAD for presenting this work at FENS 2024. We thank Björn Schwer and Cameron M. Holman for sharing our mCherry-positive ES cells. We thank Yassin Harim for computational and technical support to improve ImageJ batch workflow performance, Lorenzo Corazzi for mapping the mCherry integration position and sample preparation, Boyu Ding and the FACS core facility for cell sorting, the light microscopy core facility, the Transgenic Core facility, the Single-cell open lab, the NGS core facility, the ODCF data management core facility, and the staff at the preclinic center at the DKFZ. We also thank the imaging facility at EMBL for establishing the BrdU/EdU imaging workflow. We extend our acknowledgments to P. Ginno and T. Martin for their valuable feedback. We also thank the team members at the Wei and Höfer labs for intellectual discussions and technical support.

## Author contributions

Conceptualization: G.D.M., S.B., T.H., P.-C. W.; Study design: P.-C. W., G.D.M.; Supervision: P.-C. W., T.H.; Managing, planning, and generate chimeras: F. v.d. H., B.A.; Conducting histology, immunofluorescent experiments and imaging: G.D.M., H.-J. L., J.B.; Mathematical analysis and modeling: L.W., M.K.; Sample and library preparation: G.D.M., N.C.; Whole-genome sequencing: G.D.M., N.C.; WGS data analysis: S.B., Single-Cell RNA-sequencing analysis: L.-C. W., Visualization: G.D.M., S.B., L. W., L.-C. W., P.-C. W. and T.H.; Writing (original draft): G.D.M, S.B., L.-C. W., L.W., P.-C.W. ; Writing (review and editing): V.K., T.H..

## Data Availability

The scRNA-seq data are deposited at the GEO (GSE281055) and the WGS datasets at NLM-NCBI (Bioproject PRJNA1209637).

## Supplementary Figure Legends

**Figure S1. Characterization of tissue and cell type composition in the *NesCre::R26-DTA* chimeras. (A)** Schematic representation of the genetic strategy. Males carrying the rat *Nestin* transgenic promoter at Chr12, which drives Cre recombinase expression, were crossed with females homozygous for the *R26-DTA* transgene at Chr6, containing a Cre-dependent diphtheria toxin A (DTA) expression cassette. **(B)** H&E-stained sagittal brain sections of an *R26-DTA* and a *NesCre::R26-DTA* embryo at E12.5. Insets provided higher magnification views of rectangular areas of neocortex (ncx) or and the choroid plexus (cp). Brain regions including midbrain (mb), hindbrain (hb), diencephalon (di), neocortex (ncx), and the spinal cord (sp) were annotated. Craniofacial structures including tongue (t), and nasal cavity (n) were also indicated. Scale bars: 500 µm (overview), 50 µm (ncx view), and 20 µm (cp view). **(C)** Insets from neocortex of sagittal brain section from *R26-DTA* and *NesCre::R26-DTA* embryos at E15.5, stained against PAX6 (a marker for RG cells) and TBR2 (a marker for IP cells). Cell nuclei were counterstained with DAPI. Scale bars: 50 µm. **(D)** Bar charts showing the absolute cell counts of DAPI+, PAX6+, and TBR2+ cells in the whole brains from three *R26-DTA* and five *NesCre::R26-DTA* embryos at E15.5. Seven sagittal planes were quantified per embryo. Statistical significance was determined using an unpaired two-tailed Student’s t-test. Data are presented as mean ± SD of each sagittal plan; *p < 0.05. **(E)** A barchart showed the proportion of mCherry+ cells in the tail fibroblasts from *NesCre::R26-DTA* or *R26-DTA* chimeras at E17.5. Data were presented as mean ± standard deviation, and the statistical significance was determined by an unpaired two-tailed Student t test. n.s.: not significant. **(F,G)** Uniform Manifold Approximation and Projection (UMAP) plots showing the first and second reduced dimensions for single-cell RNA-seq datasets. Cells were visualized by brain regions **(F)**, forebrain in orange, mid-/hindbrain in blue) or by cell types **(G)** in *R26-DTA* and *NesCre::R26-DTA* chimeras, respectively.

**Figure S2. Identifying expanded subclones in the forebrain of the *NesCre::R26-DTA* chimeras. (A)** Subclone enrichment analysed by VIBER. VAF shown in Figure 2 are color- coded for the small subclones (C1, orange) and the selected/expanded subclones (C2, crimson red). Each panel represented an individual embryo. **(B)** Corrected expression level for *Nfia* (top), *Nav2* (middle), and *Ddx1* (bottom) in the cells highlighted in Figure 2C. **(C)** Schematic of the reversed blastocyst complementation strategy. *NesCre::R26-DTA* ES cells (blue) were injected into host blastocysts (DsRed) to generate *NesCre::R26-DTA* chimeras. DsRed-positive tail fibroblasts and forebrain cells were sorted by FACS at E17.5 and analysed by whole-genome sequencing, as described in Figure 2A. **(D)** VAF distribution in DsRed+ forebrain cells from three *NesCre::R26-DTA* chimeras. Each panel represented an individual mouse.

**Figure S3. Intermediate neuronal progenitors (TBR2+) display shortened cell cycles in the *NesCre::R26-DTA* chimera. (A)** EdU-BrdU pulse-chase experimental timeline. EdU was injected intraperitoneally (i.p.) at time t0, followed by BrdU at t1 (1 hour later), and embryos were collected at t2 (1.5 hours after the initial injection). **(B)** Representative immunofluorescence images of E15.5 cortical sections stained against TBR2+ (magenta) EdU (green), BrdU (red), and nuclei labelled with DAPI (grey). Images compared the isocortex region of *R26-DTA* (top) and *NesCre::R26-DTA* (bottom) chimeras. Scale bar: 50 μm. **(C)** Quantification of S-phase duration and overall cell cycle length for TBR2+ progenitors at E15.5 in *R26-DTA* and *NesCre::R26-DTA* chimeras (n = 3 embryos per group). Data were presented as mean ± SD; *p < 0.05, determined by a one-tailed unpaired Student t-test.

**Figure S4**. **Mutational signatures in *DsRed::R26-DTA* chimeras. (A)** Variant allele frequency (VAF) distributions for each substitution type (represented by different colors and stacked) in *DsRed::R26-DTA* chimeras. **(B)** Relative contribution of each substitution type across individual *DsRed::R26-DTA* chimeras.

**Figure S5. Adaptive clonal expansion altered cell type-specific pathway activity in chimera brains. (A)** Box plots showing the proportions of various cell types, including glycinergic neurons (GLYC), intermediate progenitor (IP) cells, microglia, oligodendrocyte precursor cells (OPC), pericytes, and endothelial cells, in *R26-DTA* (gray) and *NesCre::R26- DTA* (purple) chimeras. No significant differences (n.s.) were observed between groups for most cell types, except for the endothelial cells. Statistical power was determined by general linear models using Chi-square. ****: p ≦ 0.0001. **(B)** Developmental trajectory of neuroepithelial-derived cell types. Neuroepithelial cells transition to radial glial cells, which give rise to intermediate progenitors, glioblasts, oligodendrocyte precursors and ependymal cells. Intermediate progenitors further differentiate into neuroblasts and eventually generate neurons (GLYC, GABA, GLUT). Key time points (E11.5–E17.5) for differentiation were indicated. **(C–E)** Pathway-regulon activity heatmaps showing changes in the relative expression of shared genes between regulons and pathways (y-axis) and associated specific regulon activity (x-axis) in radial glial cells **(C)**, glioblasts **(D)**, and neuroblasts **(E)** in *R26-DTA* and *NesCre::R26-DTA* chimeras. Triangles represented regulons associated with the corresponding pathways (relative gene expression levels of shared gene sets) that were significantly higher (▴) or lower (▾) in their activity in *NesCre::R26-DTA* chimeras. The color of the triangles indicated statistical significance, as represented by the -log10(p) values, shown in the vertical bar at the top-right corner. The color scale at the bottom indicated regulon activity, with red representing higher activity and blue representing lower activity in *NesCre::R26-DTA* compared to *R26-DTA* chimeras.

## References

1. Chang, A. N. et al. Neural blastocyst complementation enables mouse forebrain organogenesis. Nature 563, 126–130 (2018).

2. Huang, J. et al. Generation of rat forebrain tissues in mice. Cell 187, 2129–2142.e17 (2024).

3. Stahl, R. et al. Trnp1 Regulates Expansion and Folding of the Mammalian Cerebral Cortex by Control of Radial Glial Fate. Cell 153, 535–549 (2013).

4. Fernández, V., Llinares-Benadero, C. & Borrell, V. Cerebral cortex expansion and folding: what have we learned? EMBO J. 35, 1021–1044 (2016).

5. Rakic, P. Specification of Cerebral Cortical Areas. Science 241, 170–176 (1988).

6. Rakic, P. The radial edifice of cortical architecture: From neuronal silhouettes to genetic engineering. Brain Res. Rev. 55, 204–219 (2007).

7. Borrell, V. How Cells Fold the Cerebral Cortex. J. Neurosci. 38, 776–783 (2018).

8. D’Gama, A. M. & Walsh, C. A. Somatic mosaicism and neurodevelopmental disease. Nat. Neurosci. 21, 1504–1514 (2018).

9. S., J. S., et al. Somatic Mutations in Cerebral Cortical Malformations. N. Engl. J. Med. 371, 733–743 (2014).

10. D’Gama, A. M. et al. Somatic Mutations Activating the mTOR Pathway in Dorsal Telencephalic Progenitors Cause a Continuum of Cortical Dysplasias. Cell Rep. 21, 3754– 3766 (2017).

11. Breuss, M. W. et al. Somatic mosaicism reveals clonal distributions of neocortical development. Nature 604, 689–696 (2022).

12. Körber, V., et al. Detecting and quantifying clonal selection in somatic stem cells. bioRxiv 2021.12.15.472780 (2023) doi:10.1101/2021.12.15.472780.

13. Bae, T. et al. Analysis of somatic mutations in 131 human brains reveals aging- associated hypermutability. Science 377, 511–517 (2022).

14. Gao, P. et al. Deterministic Progenitor Behavior and Unitary Production of Neurons in the Neocortex. Cell 159, 775–788 (2014).

15. Noctor, S. C., Martínez-Cerdeño, V., Ivic, L. & Kriegstein, A. R. Cortical neurons arise in symmetric and asymmetric division zones and migrate through specific phases. Nat. Neurosci. 7, 136–144 (2004).

16. Gaiano, N., Nye, J. S. & Fishell, G. Radial Glial Identity Is Promoted by Notch1 Signaling in the Murine Forebrain. Neuron 26, 395–404 (2000).

17. Nonomura, K. et al. Local Apoptosis Modulates Early Mammalian Brain Development through the Elimination of Morphogen-Producing Cells. Dev. Cell 27, 621–634 (2013).

18. Kawaue, T. et al. Lzts1 controls both neuronal delamination and outer radial glial-like cell generation during mammalian cerebral development. Nat. Commun. 10, 2780 (2019).

19. Laporte, M. H. et al. Alix is required during development for normal growth of the mouse brain. Sci. Rep. 7, 44767 (2017).

20. Ameisen, J. C. On the origin, evolution, and nature of programmed cell death: a timeline of four billion years. Cell Death Differ. 9, 367–393 (2002).

21. Yamaguchi, Y. & Miura, M. Programmed Cell Death in Neurodevelopment. Dev. Cell 32, 478–490 (2015).

22. Liang, H., Hippenmeyer, S. & Ghashghaei, H. T. A Nestin-cre transgenic mouse is insufficient for recombination in early embryonic neural progenitors. Biol. Open 1, 1200– 1203 (2012).

23. Tronche, F. et al. Disruption of the glucocorticoid receptor gene in the nervous system results in reduced anxiety. Nat. Genet. 23, 99–103 (1999).

24. Shafiee, F., Aucoin, M. G. & Jahanian-Najafabadi, A. Targeted Diphtheria Toxin-Based Therapy: A Review Article. Front. Microbiol. 10, 2340 (2019).

25. Chen, J., Lansford, R., Stewart, V., Young, F. & Alt, F. W. RAG-2-deficient blastocyst complementation: an assay of gene function in lymphocyte development. Proc. Natl. Acad. Sci. 90, 4528–4532 (1993).

26. Rivera-Pérez, J. A. & Hadjantonakis, A.-K. The Dynamics of Morphogenesis in the Early Mouse Embryo. Cold Spring Harb. Perspect. Biol. 7, a015867 (2015).

27. Junyent, S. et al. The first two blastomeres contribute unequally to the human embryo. Cell 187, 2838–2854.e17 (2024).

28. Polejaeva, I. & Mitalipov, S. Stem cell potency and the ability to contribute to chimeric organisms. Reproduction 145, R81–R88 (2013).

29. Williams, M. J. et al. Quantification of subclonal selection in cancer from bulk sequencing data. Nat. Genet. 50, 895–903 (2018).

30. Caravagna, G. et al. Subclonal reconstruction of tumors by using machine learning and population genetics. Nat. Genet. 52, 898–907 (2020).

31. Jungas, T., Joseph, M., Fawal, M.-A. & Davy, A. Population Dynamics and Neuronal Polyploidy in the Developing Neocortex. Cereb. Cortex Commun. 1, tgaa063 (2020).

32. Martynoga, B., Morrison, H., Price, D. J. & Mason, J. O. Foxg1 is required for specification of ventral telencephalon and region-specific regulation of dorsal telencephalic precursor proliferation and apoptosis. Dev. Biol. 283, 113–127 (2005).

33. Harris, L., Zalucki, O. & Piper, M. BrdU/EdU dual labeling to determine the cell-cycle dynamics of defined cellular subpopulations. J. Mol. Histol. 49, 229–234 (2018).

34. Alexandrov, L. B. et al. The repertoire of mutational signatures in human cancer. Nature 578, 94–101 (2020).

35. Silbereis, J. C., Pochareddy, S., Zhu, Y., Li, M. & Sestan, N. The Cellular and Molecular Landscapes of the Developing Human Central Nervous System. Neuron 89, 248–268 (2016).

36. Zheng, X. et al. Metabolic reprogramming during neuronal differentiation from aerobic glycolysis to neuronal oxidative phosphorylation. eLife 5, e13374 (2016).

37. Ginhoux, F. & Prinz, M. Origin of Microglia: Current Concepts and Past Controversies. Cold Spring Harb. Perspect. Biol. 7, a020537 (2015).

38. Lendahl, U., Zimmerman, L. B. & McKay, R. D. G. CNS stem cells express a new class of intermediate filament protein. Cell 60, 585–595 (1990).

39. Morshead, C. M. et al. Neural stem cells in the adult mammalian forebrain: A relatively quiescent subpopulation of subependymal cells. Neuron 13, 1071–1082 (1994).

40. Kawaguchi, A. et al. Single-cell gene profiling defines differential progenitor subclasses in mammalian neurogenesis. Development 135, 3113–3124 (2008).

41. Chapman, M. S. et al. Clonal dynamics after allogeneic haematopoietic cell transplantation. Nature 635, 926–934 (2024).

42. Dai, H.-Q. et al. Direct analysis of brain phenotypes via neural blastocyst complementation. Nat. Protoc. 15, 3154–3181 (2020).

43. Pachitariu, M. & Stringer, C. Cellpose 2.0: how to train your own model. Nat. Methods 19, 1634–1641 (2022).

44. Stringer, C., Wang, T., Michaelos, M. & Pachitariu, M. Cellpose: a generalist algorithm for cellular segmentation. Nat. Methods 18, 100–106 (2021).

45. Khan, M. & Gasser, S. Generating Primary Fibroblast Cultures from Mouse Ear and Tail Tissues. J. Vis. Exp. : JoVE (2016) doi:10.3791/53565.

46. Islam, S. M. A. et al. Uncovering novel mutational signatures by de novo extraction with SigProfilerExtractor. Cell Genom. 2, 100179 (2022).

47. Wolock, S. L., Lopez, R. & Klein, A. M. Scrublet: Computational Identification of Cell Doublets in Single-Cell Transcriptomic Data. Cell Syst. 8, 281–291.e9 (2019).

48. Hao, Y. et al. Integrated analysis of multimodal single-cell data. Cell 184, 3573–3587.e29 (2021).

49. Korsunsky, I. et al. Fast, sensitive and accurate integration of single-cell data with Harmony. Nat. Methods 16, 1289–1296 (2019).

50. Manno, G. L. et al. Molecular architecture of the developing mouse brain. Nature 596, 92–96 (2021).

51. Sande, B. V. de et al. A scalable SCENIC workflow for single-cell gene regulatory network analysis. Nat. Protoc. 15, 2247–2276 (2020).

52. Aibar, S. et al. SCENIC: Single-cell regulatory network inference and clustering. Nat. methods 14, 1083–1086 (2017).

53. Korotkevich, G. et al. Fast gene set enrichment analysis. bioRxiv 060012 (2021) doi:10.1101/060012.

54. Liberzon, A. et al. The Molecular Signatures Database Hallmark Gene Set Collection. Cell Syst. 1, 417–425 (2015).

