## Supplementary figures and images for "Adaptive Clonal Expansion Shapes Brain Development"

### Figure S1

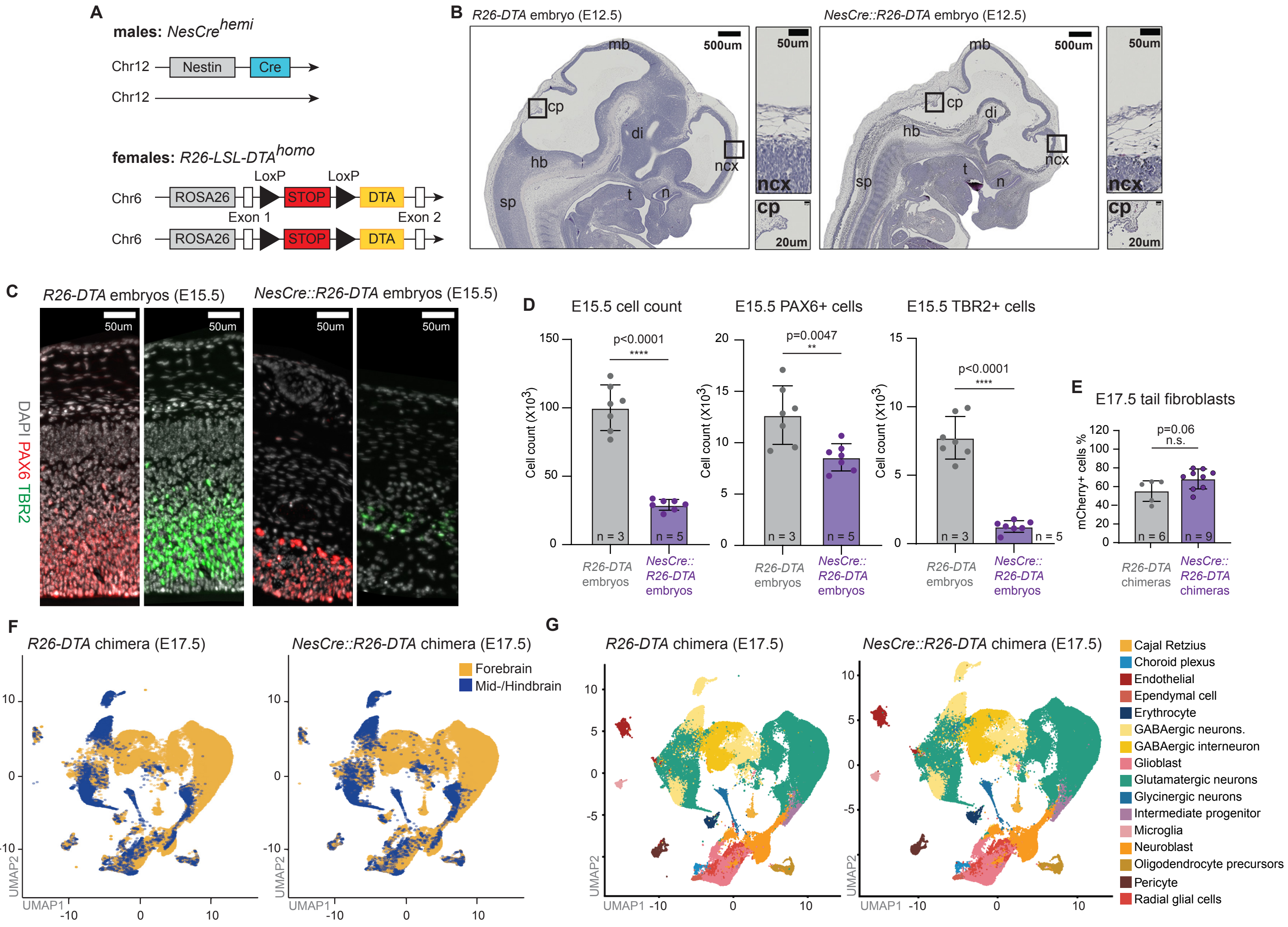

### Figure S2

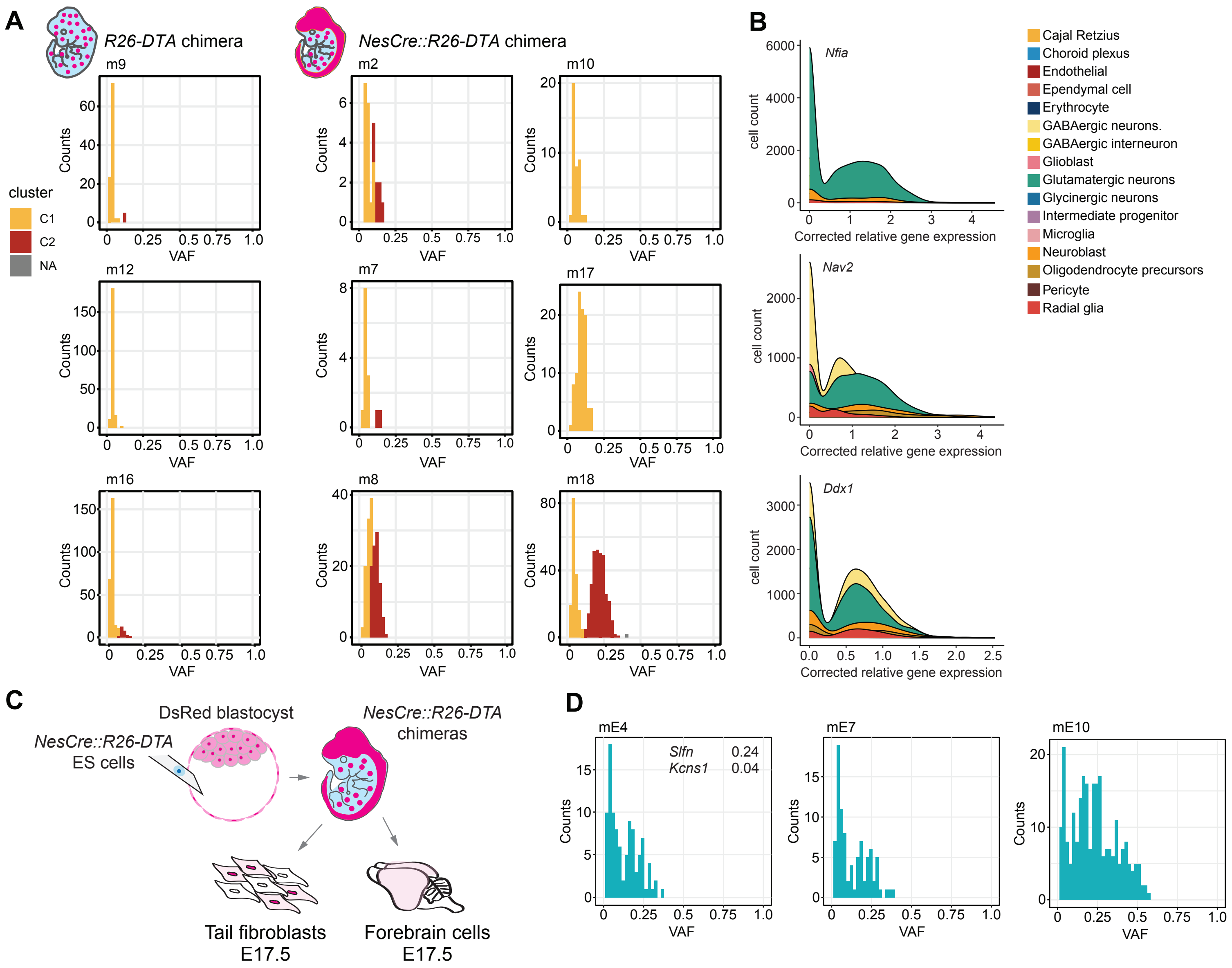

### Figure S3

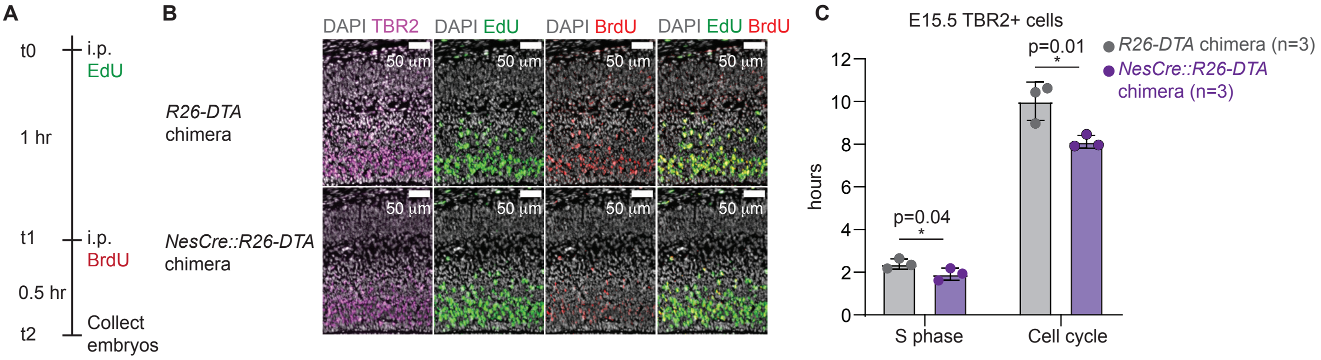

### Figure S4

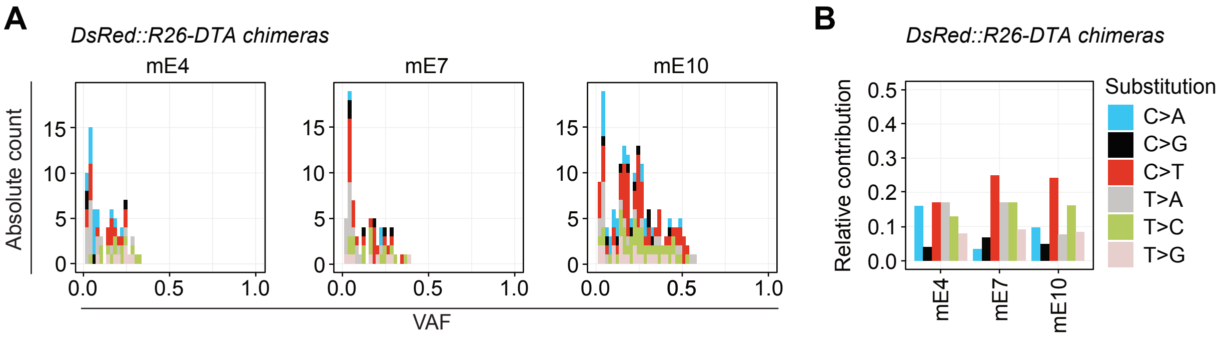

### Figure S5

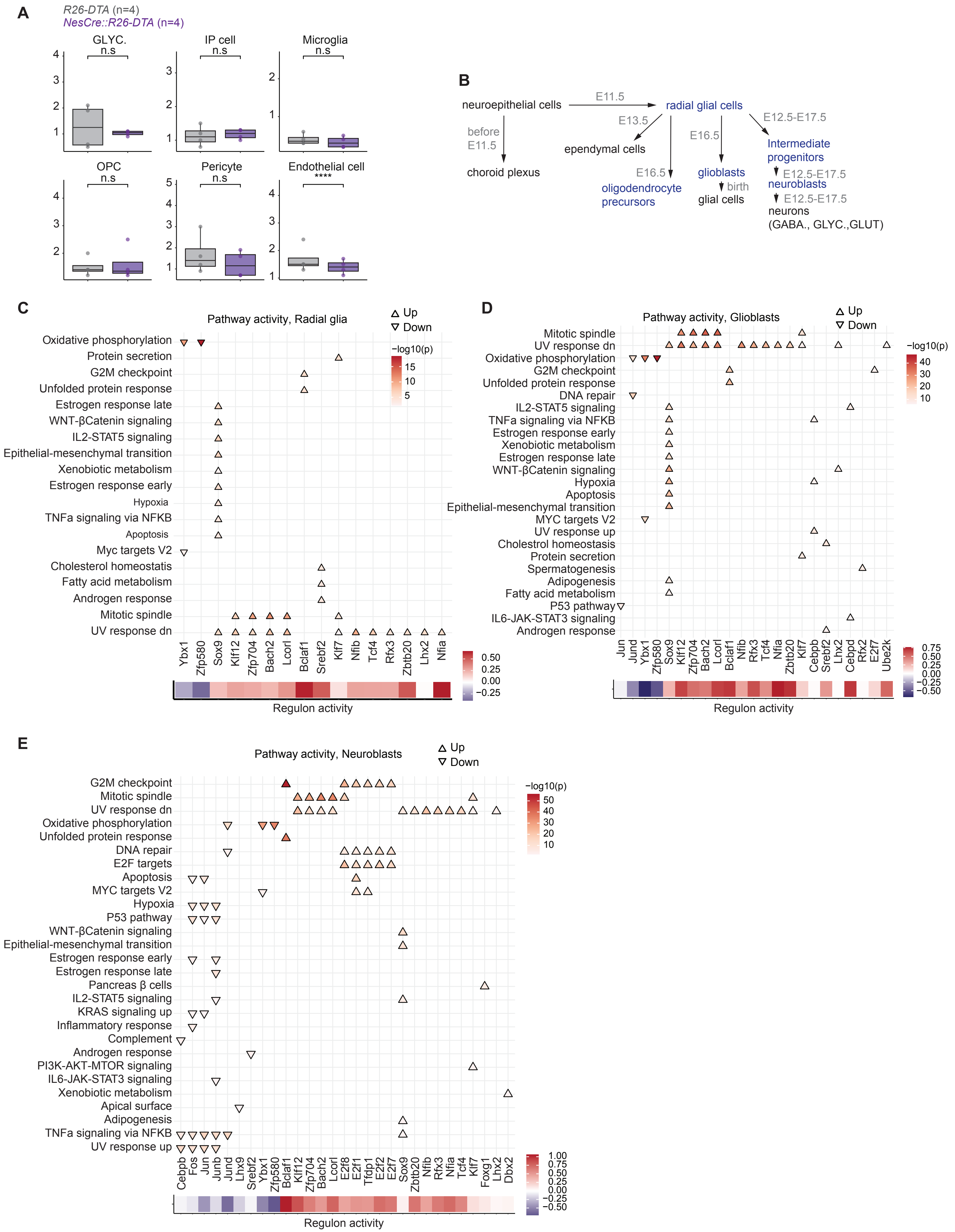
